# Role of HAO2 in rats with chronic kidney disease by regulating fatty acid metabolic processes in renal tissue

**DOI:** 10.1101/2022.12.13.520357

**Authors:** Xunjia Li, Chengxuan Liu, Aimin Yang, Youfeng Shen, Jian Xu, Deyu Zuo

## Abstract

Fibrosis is a progressive, often irreversible histologic manifestation of chronic and end-stage renal disease. In this study, single-cell transcriptome sequencing technology was used to sequence and analyze blood and kidney tissue cells in normal control rats and rats with chronic kidney disease (CKD), focusing on key cell populations and functional enrichment to explore the pathogenesis of CKD. Oil red O staining and ELISA were used to detect lipid droplets and free fat acid (FFA). RT-PCR, WB were used to verify the differential gene HAO2 and fatty acid metabolic process in tissue to ensure the reliability of single-cell sequencing results. We successfully established a single-cell transcriptome atlas of blood and kidney tissue in rats with CKD, which were annotated into 14 cell subsets (MPCs, PT, Tc, DCT, B-IC, A-IC, CNT, ALOH, BC, Neu, Endo, Pla, NKT, Baso) according to marker gene, and the integrated single-cell atlas of rats showed a significant increase and decrease of MPCs and PTs in the model group, respectively. Functional analysis found extensive enrichment of metabolic-related pathways in PT cells, includes fatty acid metabolic process, cellular amino acid metabolic process and generation of precursor metabolites and energy. Immunohistochemical experiments determined that the differential gene HAO2 was localized in the renal tubules, and its expression was significantly reduced in model group compared with control, and oil red O staining showed that lipid droplets increased in the model group. ELISA assay showed that ATP content decreased in the model group and FFA increased in the model group. ACOX1, PPARα, PGC1α were decreased in the model group, while genes and proteins were increased after overexpression of HAO2, and the AMPK and ACC phosphorylated proteins were increased. Therefore, HAO2 may be an important regulator of fatty acid metabolic processes in CKD, and overexpression of HAO2 can enhance fatty acid metabolism by promoting fatty acid oxidation pathway.

## Introduction

Chronic kidney disease (CKD) refers to a disease characterized by irreversible damage to the kidneys (≧ 3 months) caused by multiple etiologies (chronic nephritis, diabetes, hypertension, etc.) [1]. There are five subtypes of CKD classified by causes which include type 1 diabetes, type 2 diabetes, hypertension, glomerulonephritis, and other causes [2]. CKD is identified by a decrease in glomerular filtration rate (eGFR) or signs of kidney damage with abnormal pathological, blood, urine, or imaging tests. And the features of renal damage are glomerulosclerosis, tubular atrophy, and interstitial fibrosis.

Generally, the glomerular filtration rate is between 30-89 mL/min per 1.73 m^2^, which belongs to the stage 2∼3 of CKD [3]. The treatment of CKD is mainly to delay the progression of the disease and reduce complications, with the gradual decline of glomerular filtration rate, the decline of renal function, the accumulation of toxins, nephropathy develops to CKD stage 5, that is, end-stage renal disease (ESRD), commonly known as uremia [4, 5]. Around 10% of the global population was affected by CKD, CKD has become a global public health problem [6]. Approximately 120 million adult patients in China, accounting for 10.8 percent of the national population, will eventually progress to ESRD and can only be sustained by kidney dialysis or kidney transplantation [7]. Therefore, early CKD intervention protection is particularly important to avoid serious disease progress.

At present, the pathological mechanism of CKD has not been fully elucidated, but one view is that oxidative stress is involved in CKD progression. Low concentrations of ROS are thought to regulate signaling pathways in the kidneys related to mitochondrial function, which is closely related to renal homeostasis maintenance [8, 9]. The mechanism of renal disease is related to multiple factors, including autophagy, inflammation, mitochondrial dysfunction, and oxidative stress, especially autophagy activation has been implicated in clinical studies such as CKD and acute kidney injury[10]. In addition, studies have shown that gut microbial dysregulation is also related to the pathogenesis of CKD [11]. Single-cell RNA sequencing (scRNA-seq) has played an increasingly important role in biological research, offering unique and significant advantages in revealing cell-to-cell heterogeneity in tissue blocks (or cell suspensions) and resolving the functions and characteristics of individual cells. Therefore, it has been widely used in cell biology, immunology and oncology in recent years.

In this study, scRNA-seq was carried out by constructing a rat model of CKD, and single-cell atlas of rat with CKD were drawn, aiming to find out the important cell subsets and pathway enrichment in the process of CKD, and verify the results by designing *in vitro* validation experiments. It is hoped that these findings will provide evidence of functional heterogeneity of important cell clusters at the single-cell level and the pathogenesis of differential genes in CKD.

## Materials and methods

### CKD model in rats

After 7 days of adaptive rearing, SD rats were divided into 2 groups (control group and CKD group), 4 in each group, 3 for scRNA-seq sampling, and 1 in reserve. Reference of CKD model construction and self-exploration to finally determine the model construction method [12]. The model group was continuously gavaged with 0.25% adenine (McLean, catalog number: A800684) suspension for 8 weeks (150mg/kg/d), and the control group was gavaged with the same volume of distilled water. The rat was anesthetized and sacrificed, blood was taken from the abdominal cavity for backup, and the kidney tissue was taken in GEXSCOPE® tissue preservation solution for storage for backup.

### Single-cell RNA sequencing

The tissue sample was washed 3 times with Hanks Balanced Salt Solution (HBSS) and cut into 1-2 mm pieces. Then add 2 ml GEXSCOPE® tissue dissociation solution and digest the tissue blocks in a 37 °C, 15 ml centrifuge tube with continuous agitation for 15 min. After digestion, filter the samples with a 40 µm sterile strainer and centrifuge at 1,000 rpm for 5 min. Then discard the supernatant and resuspend the pellet with 1 ml PBS. Add 2 ml of GEXSCOPE erythrocyte® lysis buffer for 10 min at 25 °C to remove erythrocytes. Centrifuge at 500 rpm for 5 min and suspend in PBS. Stain with trypan blue (Sigma) and evaluate under a microscope. Prepare a phosphate buffer single-cell suspension at a concentration of 1×10 ^5^ cells/mL, load the single-cell suspension onto a microfluidic device, and construct a scRNA-seq library according to the Singleron GEXSCOPE protocol (Singleron Biotechnologies). Once the library is constructed, dilute to 4 nM and sequence with 150 bp paired end reads on the IlluminaNovaSeq6000. CeleScope (http://github.com/singleron-RD/CeleScope), was used to process the raw readings (reads) from the disembarking machine to generate an RNA expression matrix.

### Quality control, dimensionality reduction, and cluster analysis

Cells were filtered with a gene count between 200 and 1500 and a UMI count below 30,000, removing cells with a mitochondrial content of more than 45% and cells with a red blood cell content of more than 0.025%, and a total of 27,332 cells were obtained after filtration. The functions in the Seuratv4.1.1 program were used for dimensionality reduction and clustering, and the harmony algorithm performed batch correction on the gene expression matrix. NormalizeData() and ScaleData() normalize all gene expressions. FindVariableFeatures() selects the first 2000 variable genes for PCA analysis. FindCluster() uses the first 30 principal components with the resolution parameter set to 0.5 to cluster the cells. The tSNE algorithm is used for two-dimensional spatial cell cluster visualization.

### Differential genetic analysis

In this study, the differential expression analysis of mononuclear phagocytic cells (MPCs) and proximal tubular (PT) between the control group and the model group was mainly performed. Seurat v4.1.1 FindMarkers() was based on the Wilcox statistical method with default parameters and filter genes as differentially expressed genes (DEGs) which is expressed in more than 10% of cells in the cluster and |avg_log2FC| > 0.5, p_val < 0.05.

### Function enrichment analysis

GO and KEGG enrichment analyses were performed together using the ClusterProfiler R package. The pathway whose p_adj < 0.05 is considered to be significantly enriched.

### Immunohistochemistry

The kidney tissue of rats was taken for paraffin embedding, dewaxed and hydrated by xylene and ethanol solution of different concentrations, and antigen repair was carried out with immunohistochemical repair solution with pH 8 under high temperature and pressure, and after repair, it was cleaned with tap water and blocked with catalase solution, and the primary antibody diluent was incubated after blocking, and the secondary antibody working solution was incubated at room temperature for 20 minutes after washing with tap water. Incubate the enzyme-labeling solution for 20 min after washing again, and use DAB chromogenic solution for color development after washing. Then hematoxylin staining, washing, hydration, transparent were performed sequentially and finally sealed.

### Oil red O staining

The tissues were fixed with 4% paraformaldehyde for 10 min. After washing, add 1-2 mL of oil red O working solution, stain at room temperature in the dark for 30 min. Rinse with 60% isopropyl alcohol for 30s until the background is transparent and continue to wash with water. Hematoxylin staining solution counterstained nuclei for 30s, washed with water until bluish, sealed with glycerol gelatin. After drying at room temperature, microscopic observation of lipid droplet staining was conducted.

### ELISA assay

The Rat ATP ELISA Kit (m1460314) and FFA ELISA Kit (ml092767) are products of Shanghai Enzyme-linked Biotechnology Co., Ltd. Extract rat whole blood according to the instructions of ELISA kit, centrifuge and then take the supernatant for later use; Add standards, samples, enzyme reagents according to the instructions, wash after incubation, and then add color developer and stop solution in turn, use a multifunctional microplate reader to determine the absorbance of each well at a wavelength of 450 nm within 5 minutes, calculate the regression equation of the standard curve with the concentration and absorbance value of the standard, and then calculate the sample concentration.

### qPCR

Kidney tissue was extracted, an appropriate amount of TRIzol (1 mL/50-100 mg of kidney tissue) was added, the supernatant was centrifuged after grinding, chloroform (200 μL/1 mL TRIzol) was added, the supernatant was centrifuged after shaking and mixed for 15 min, and isopropanol (0. 5 mL/1 mL TRIzol) was added, shake and mix well for 10 min, centrifuge and discard the supernatant, add an appropriate amount of 75% ethanol (1 mL/1 mL TRIzol), shake the suspension precipitate, centrifuge and discard the supernatant, add 50 μL of RNA-free enzyme, water shaking and mix, and test the RNA concentration on the machine. The reverse transcription kit reagent was added, the RNA was reverse-transcribed into complementary DNA by amplification, primers and SYBR were added, and the Ct values of AMPK, ACC, ACOX1, PPARα and PGC1α were detected by a real-time PCR instrument, and the results were expressed as 2^−ΔΔCt^. The PCR-primers used for PCR are as follows: AMPK-F 5’-GGCAAGCACTTCATCAACCG-3’, AMPK-R 5’-GCACACGGAAAAGATCAGCG-3’; ACC-F 5’-TCTGGGACGTCGAAATTGCC-3’, ACC-R 5’-

GTTTTGGCCAACGGAGATGG-3’; ACOX1-F 5’-

CTGGAGTTTAGGGGAAGCCG-3’, ACOX1-R 5’-

GCCTTTAAACGCCATGCAAGA-3’; PPARα-F 5’-

CCTTACCCCCAGCGTTTGTT-3’, PPARα-R 5’-

CAAGCTGAGTGGAGCATGAG-3’; PGC1α-F 5’-

TCCTCATTGGTTGACGGAGC-3’, PGC1α-R 5’-

TCATCCACCTGACTGTTGCC-3’.

### Western blotting

Extract each group of tissue proteins according to the instructions of the protein extraction kit, use BCA method for protein quantification, add 5 × protein loading buffer after aliquoting, boil in a 100 °C water bath for 10 min, and store at −80 °C for later use. The gel was prepared according to the sodium lauryl sulfate-polyacrylamide gel electrophoresis kit, and after loading, electrophoresis and transfer, the primary antibody diluted at 1:1000 (AMPK, abclonal, A12491; p-AMPK, abclonal, AP1002; ACC, abclonal, A19627; p-ACC, abclonal, AP0298; ACOX1, abclonal, A21217; PPARα, abclonal, A18252; PGC1α, abclonal, A12348; Hao2, abclonal, A15159; GAPDH, abclonal, A19056) was incubated overnight and the secondary antibody (1:2000, abclonal, AS014) was incubated for 1h. The band was exposed by the gel imaging instrument, and the grayscale value of the band was analyzed by Image J software.

## Results

### Single-cell atlas of rats with CKD

SD rats were constructed using low-dose adenine to induce CKD, and a rat control group was set up. Two groups of rat blood samples and kidney tissue samples were taken for scRNA-seq (D: control; M: Model; D/M-number (7,8,9,10,11,4) represents the blood sample; D/M-letter (H,I,J,D,L,M) represents the kidney tissue specimen). After quality control, a total of 27,332 cells were obtained, the retained genes were pretreated, and the first 2000 variable genes were selected for PCA dimensionality reduction, and finally the first 30 PCs were selected for subsequent cluster analysis and visualization. As shown in Figure 1A, cells are clustered into 23 cell subsets that are annotated into 14 cell subsets (MPCs, PT, Tc, DCT, B-IC, A-IC, CNT, ALOH, BC, Neu, Endo, Pla, NKT, Baso) according to the marker gene (Figure 1B). The t-SNE visualization between groups showed that MPCs, Neu increased in the model group and PT cells, Tc, B-IC, and A-ICs decreased in the model group (Figure 1C, 1G). The cell annotated marker genes are Cd68, C1qa, Lyz2, Cd3g, Trbc2, Slc12a3, Slc26a4, Slc4a1, Mme, Calb1, Umod, Cd79b, S100a8, Lcn2, Gatm, Emcn, J chain, Col3a1, Mcpt1l4, and its relative expression in cell populations and cell types is presented using violin diagrams, heatmaps and scatterplots (Figures 1D-F). The distribution of the contribution of cells derived from different rat blood samples or kidney tissue to cell clustering and typing is shown in Figure 1H.

**Fig. 1.**
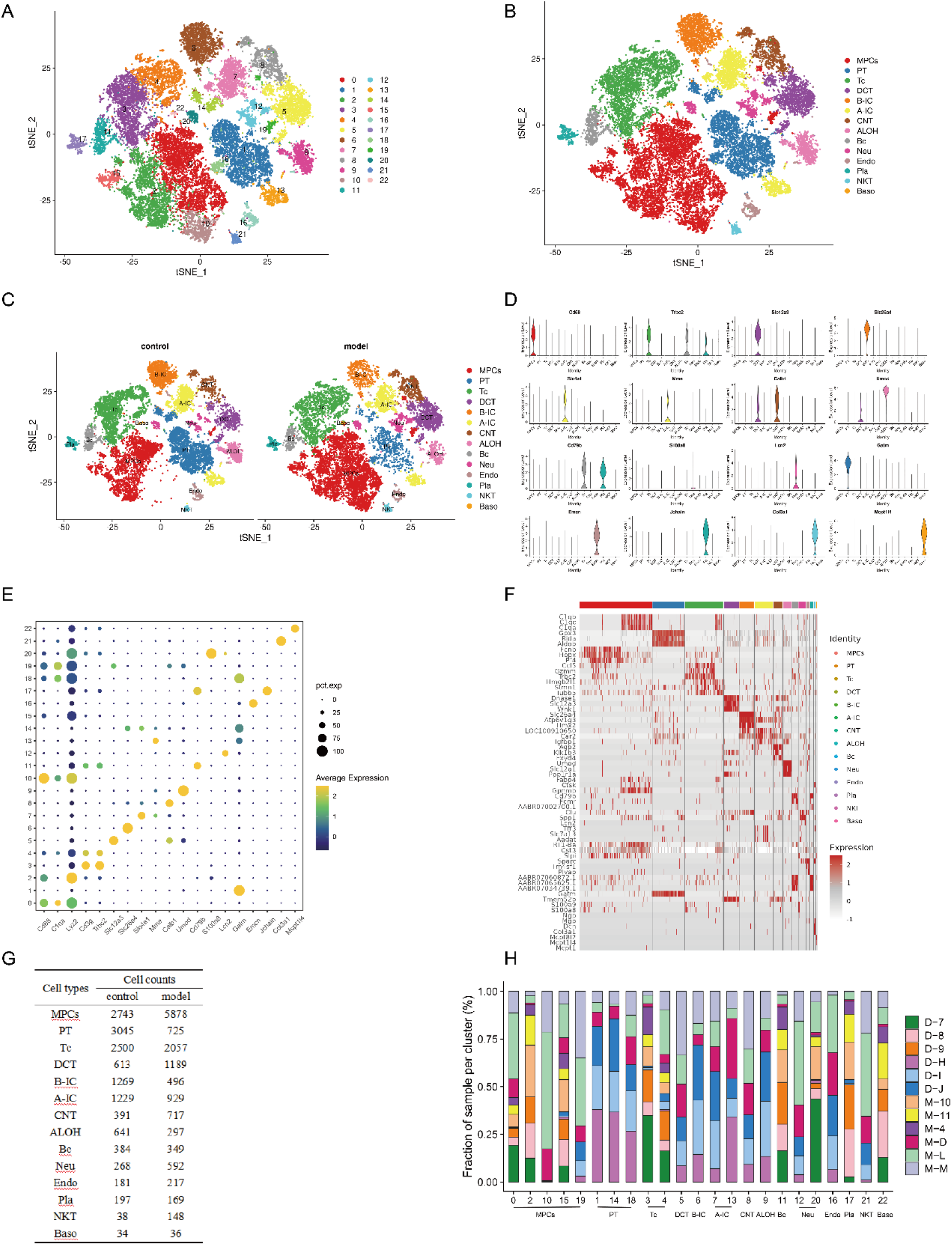
Single-cell transcriptome analysis of rat blood and kidney tissue. A, t-SNE visualization of cell clustering. B, t-SNE visualization of cell type. C, t-SNE visualization plots of cell type in the control and model groups. D, Violin diagram of differential genes in different cell types. E, Scattered distribution of differential genes in different cell clusters. F, The heatmap of marker gene in cell typing. G, Table shows the quantitative distribution of cell types in different groups. H, The distribution plot of the contribution of cells to cell type in different samples. D: control; M: Model; The D/M-number (7,8,9,10,11,4) represents the blood sample; The D/M-letter (H,I,J,D,L,M) represents the kidney tissue specimen Abbreviations: MPCs, mononuclear phagocytic cell; PT, proximal tubule; Tc, T cell; DCT, distal convoluted tubule; B-IC, beta intercalated cell; A-IC, alpha intercalated cell; CNT, connecting tubule; ALOH, ascending loop of Henle; Bc, B cell; Neu, neutrophil; Endo, endothelial cell; Pla, plasma cell; NKT, natural killer cell; Baso, basophile

### Analysis of the evolution of cells in rat blood and renal tissue specimen

To further analyze the evolution of cells from rat blood samples and kidney tissue, we performed a pseudotime analysis with Monocle 3.0. As shown in Figure 2A, the image on the left is the timeline of cell development, the darker the color represents the earlier the development stage, the image on the right is the developmental stage and specific information of each subpopulation of cells, the results show that MPCs, Tc, Pla are mainly in the early stage of development, Endo cells are in the early stage of development, most of them are in the late evolution, and PT cells increase significantly in the late evolution. Combined with the single-cell atlas of rats in Figure 1, it was suggested that in the process of transforming rats from normal to CKD, the trajectory of cell evolution, PT and Endo were at the beginning of differentiation, and MPCs and NKT increased significantly with the disease process. When processed in state2 and state3, the relative expression of Atp5i was significantly increased, which was consistent with the differentiation trajectories of Endo and PT cells, and Gatm increased significantly in the late stage of the pseudotime sequence, which was consistent with the differentiation trajectories of PT cells (Figure 2B-D).

**Fig. 2.**
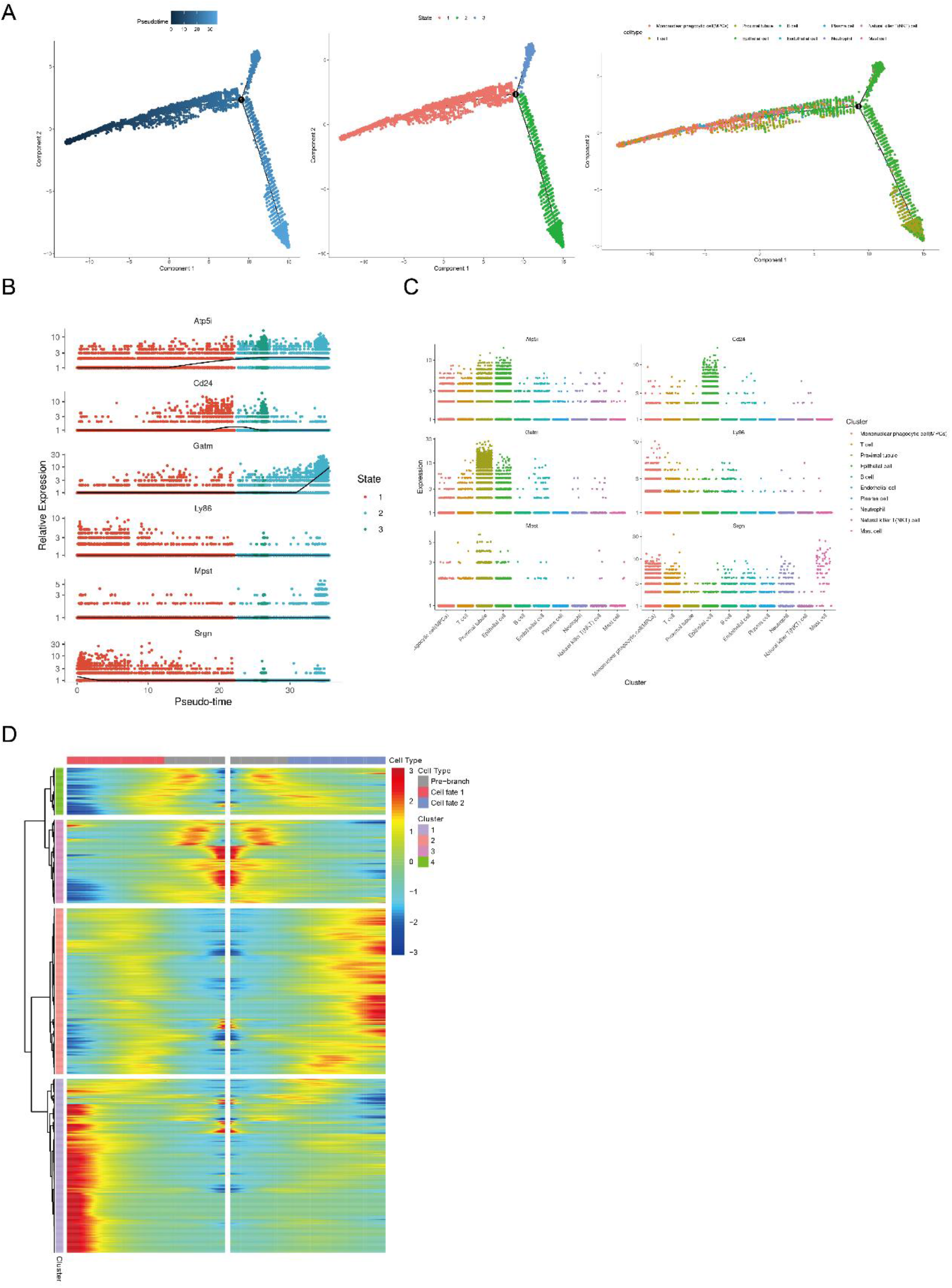
Differentiation trajectories of cell subsets of rat blood and kidney tissue. A, The reference timeline of cell development and the developmental stage of the cell subset. B, The relative expression of genes in cell development trajectories. C, The relative genes expression of different cell subsets in developmental trajectories. D, Gene expression heatmap of cell developmental trajectories.

### Clustering of MPCs subsets

The rat-integrated single-cell atlas showed a significant increase in MPCs in the model group, while pseudotime analysis showed that cells differentiated along MPCs trajectories with disease progression. MPCs cells were extracted for further analysis, and found MPCs cells were divided into 15 cell subsets (Figure 3A) and cell-annotated separately (Figure 3B). 15 cell subsets were annotated with 15 cell types according to the marker gene: C0-Monocyte1, C1-Macrophages1, C2-Monocyte2, C3-Plasmacytoiddendritic, C4-Macrophages2, C5-Tcell1, C6-Monocyte3, C7-Macrophages3, C8-Plasmacytoiddendritic, C9-Macrophages4, C10-Monocyte4, C11-Macrophages5, C12-Unknown, C13-Macrophages6, C14-Tcell2. The t-SNE visualization showed the distribution of MPCs subsets in the control group and the model group, and macrophages such as C1, C4, C7, C9, C11, C13 and monocytes such as C2, C6, C10 increased significantly in the model group (Figure 3C, 3G). The relative expression of cell-annotated marker genes in cell populations and cell types is presented using violin plots, heatmaps, and scatter plots (Figure 3D-F). The contribution distribution of cells derived from different rat blood samples or kidney tissue to the cell typing of MPCs is shown in Figure 3H.

**Fig. 3.**
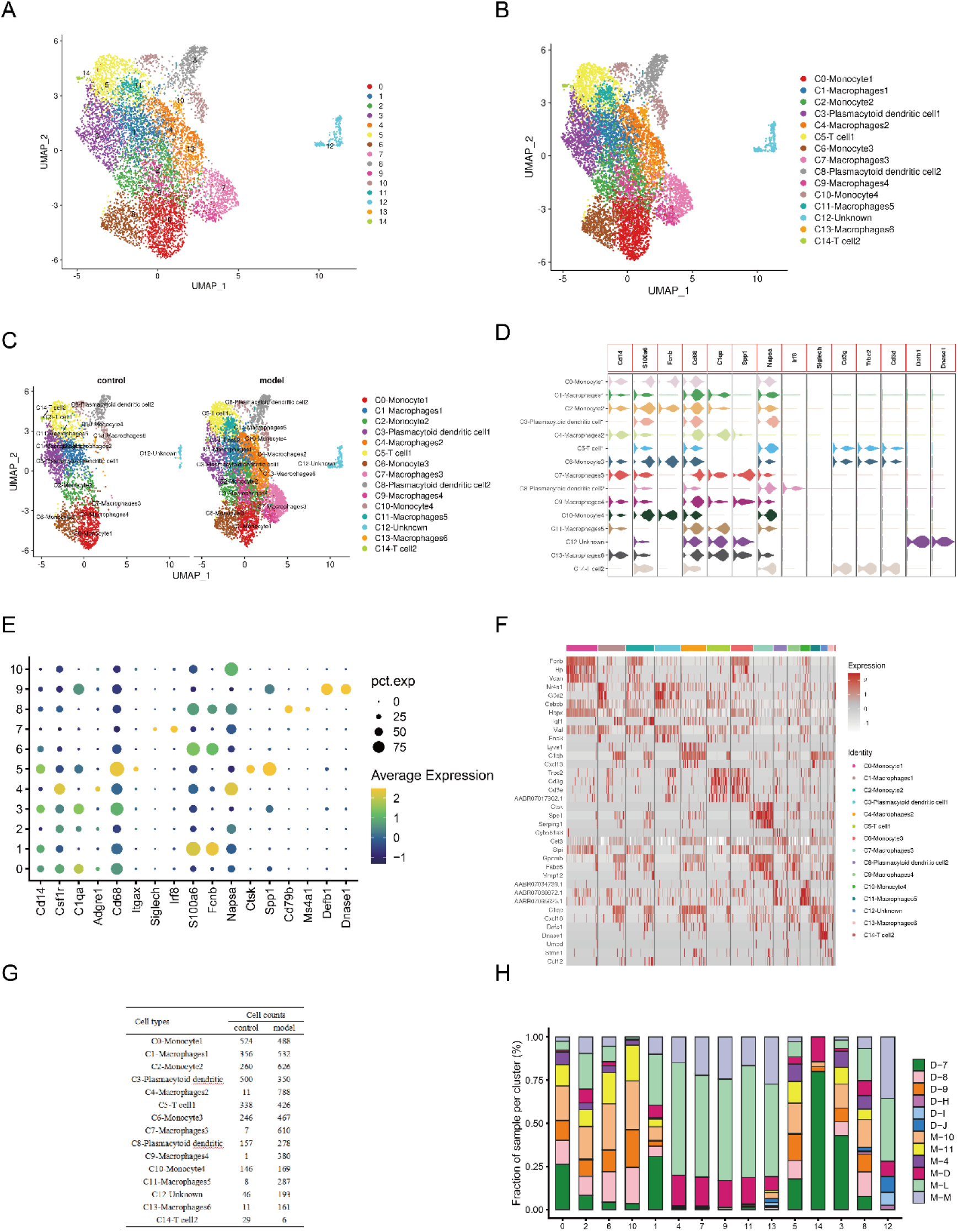
Single cell atlas of MPCs. A, Re-clustering of MPCs using UMAP algorithm. B, UMAP visualization of MPCs cellular heterogeneity. C, UMAP visualization of heterogeneity of MPCs cells in the control and model groups. D, The violin diagram of marker gene. E, Scatter distribution of marker genes in MPCs re-clustered cell clusters. F, The heatmap of marker genes in MPCs re-clustered cell clusters. G, Table shows the quantitative distribution of different subtypes of MPCs in the control and model group. H, The distribution plot of the contribution of cells to different subtypes of MPCs in different samples.

Similarly, we used Monocle 3.0 software to further analyze the evolution of MPCs cells. As shown in Figure 4A, MPCs cell development is divided into 7 states, C3 is mainly in state 1, C8 is in state 7 and state 5; C6 is mainly in state 1, state 2 and state 7, while C10 is in state 1 and state 3. The relative expressions of Anxa3, Cstb, ler3, Ptprc, RT1-Ba and Srgn in MPCs at different developmental stages and in different cell types are shown in Figure 4B-C. The relative expression heatmap of genes of cell fate branches is shown in Figure 4D-E.

**Fig. 4.**
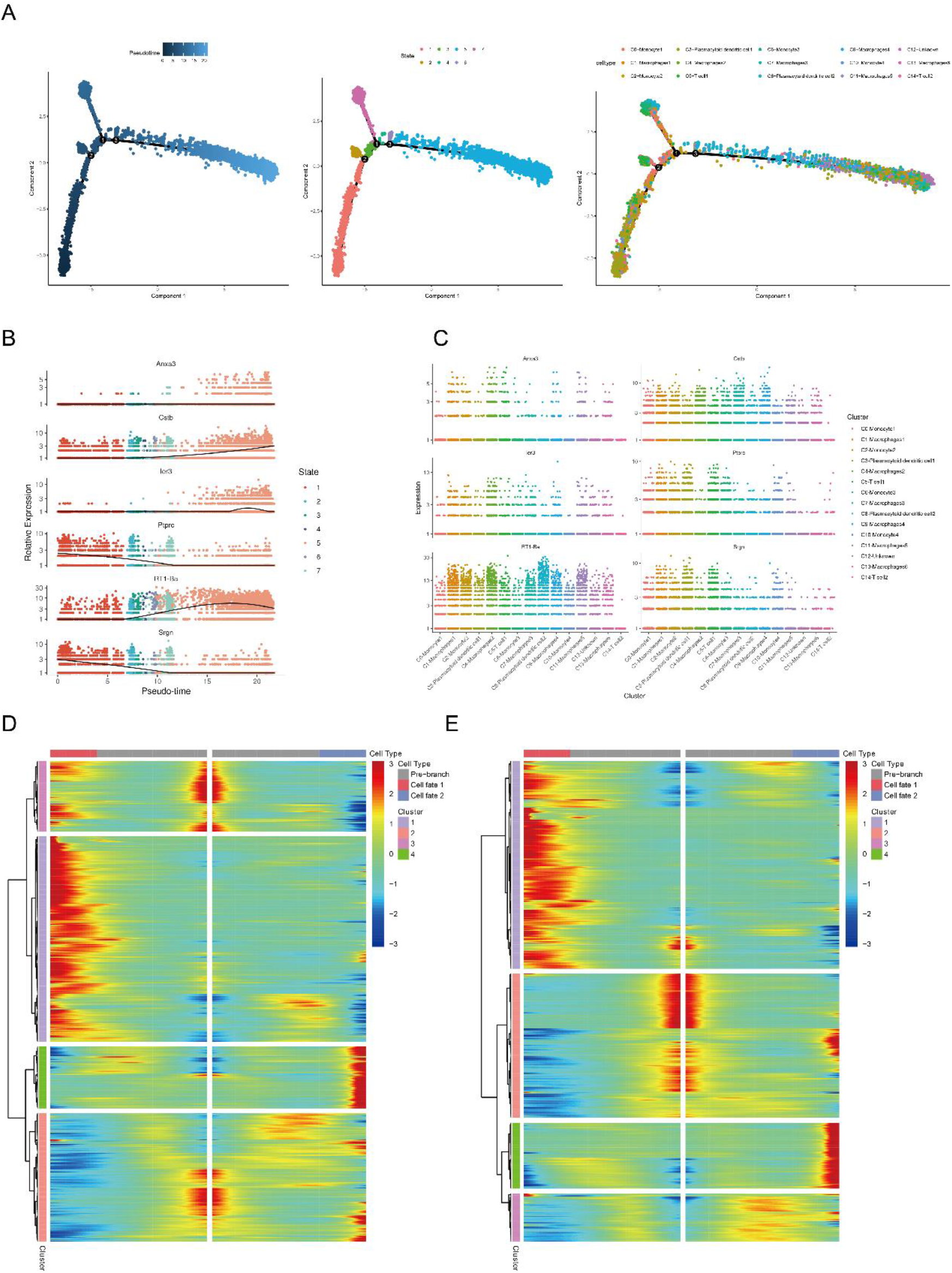
Differentiation trajectories of subgroups of MPCs re-clustering. A, The reference timeline for MPCs cell development. B, The relative expression of genes in the developmental trajectories of MPCs cells. C, The relative genes expression of different MPCs cell subsets in developmental trajectories. D, Gene expression heatmap of cell branch 1. E, Gene expression heatmap of cell branch 2.

### Functional enrichment analysis of MPCs

The differential expression and enrichment analysis of genes in the control and model groups in the MPCs cell population are shown in Figure 5. Compared with the control group, a total of 160 genes were upregulated, and 196 genes were downregulated in the model group (Figure 5A). GO enrichment function analysis showed that the upregulated gene in the control group was mainly related to leucocyte cell-cell adhesion, phagocytosis, positive regulation of cytokine production, etc., and the cell components (CC) involved were external side of plasma membrane, actin Cytoskeleton is dominant (Figure 5B). KEGG pathway analysis showed that the upregulated genes in the control group were mainly related to the Epstein-Barr virus infection, Cell adhesion molecules, and Regulation of actin cytoskeleton pathways (Figure 5C). The downregulated genes are mainly biological processes (BP) such as negative regulation of immune system process, ATP metabolic process and response to glucocorticoid (Figure 5D), while KEGG pathway is mainly enriched in Lysosome, Rheumatoid arthritis, Chemical Carcinogenesis-reactive oxygen species, Diabetic cardiomyopathy and Oxidative phosphorylation, which are also associated with the PPAR signaling pathway (Figure 5E).

**Fig. 5.**
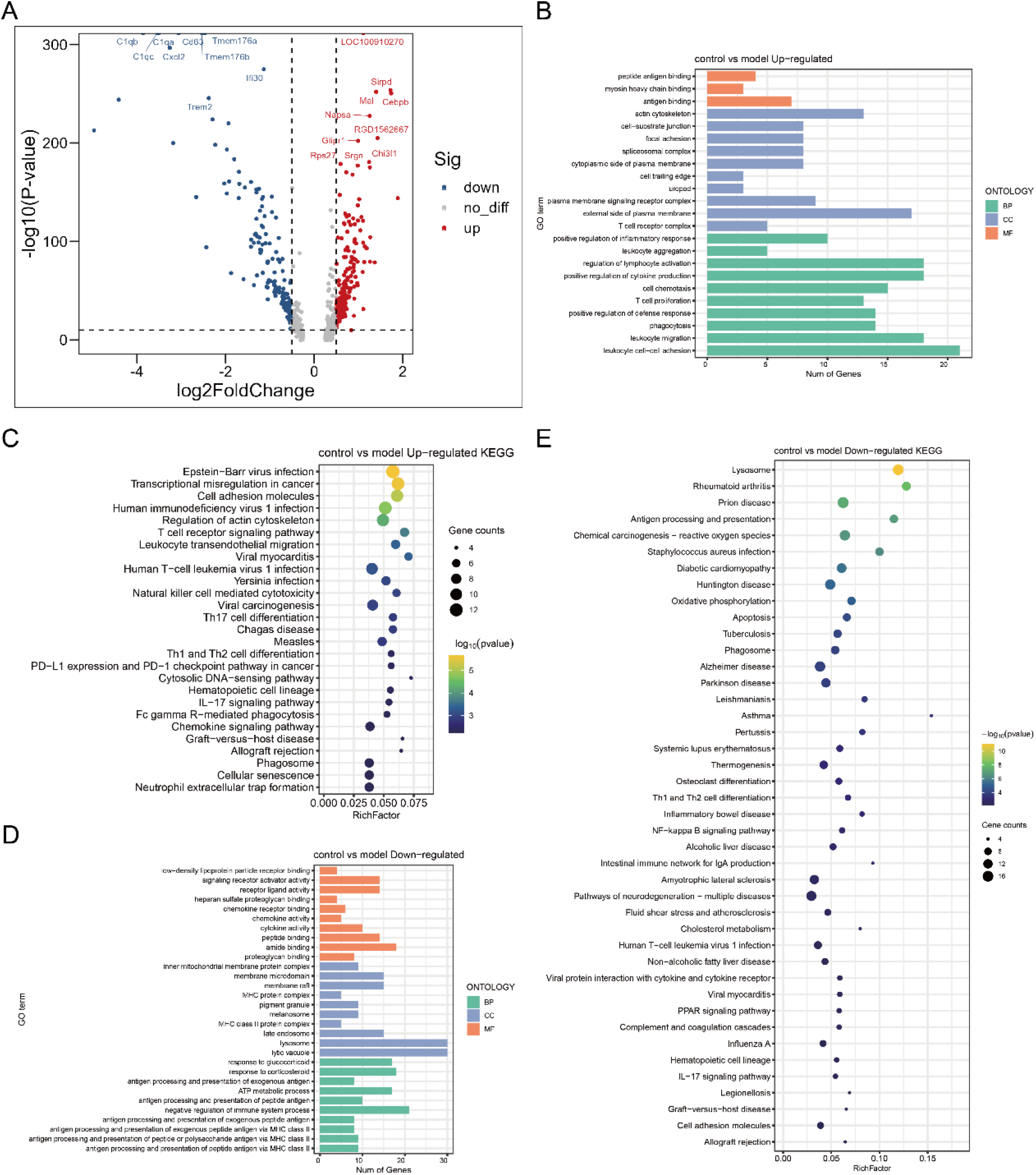
After the MPCs were re-clustered, the differential gene enrichment analysis between the control and model group was analyzed. A, Volcano plot of differential genes between control and model groups. B, GO enrichment of differential genes upregulated by the control group compared to the model group. C, Enrichment of KEGG pathway of differential genes upregulated in the control group compared with the model group. D, GO enrichment of differential genes downregulated by the control group compared with the model group. E, Enrichment of KEGG pathway of differential genes downregulated by the control group compared with the model group.

### Analysis of PT cell subset

PT cells were extracted for further analysis, and were clustered into 10 cell subsets (Figure 6A) and cell-annotated separately, and the 10 cell subsets were annotated into 3 cell types according to the marker gene: Proximal_tubule1 (PCT), Proximal_tubule2 (PST), and Macrophages (Figure 6B). Proximal_tubule1, Proximal_tubule2 was significantly reduced in the model group (Figure 6C, 6G). Violin plots, heatmaps, and scatterplots are used to show the distribution of relative expression of the marker gene in subpopulations and classifications of PT cell typing (Figure 6D-F). The contribution of cells derived from different rat kidney tissues to PT cell typing is shown in Figure 6H, and it is obvious that PT cells are mainly derived from kidney tissue of the control group.

**Fig. 6.**
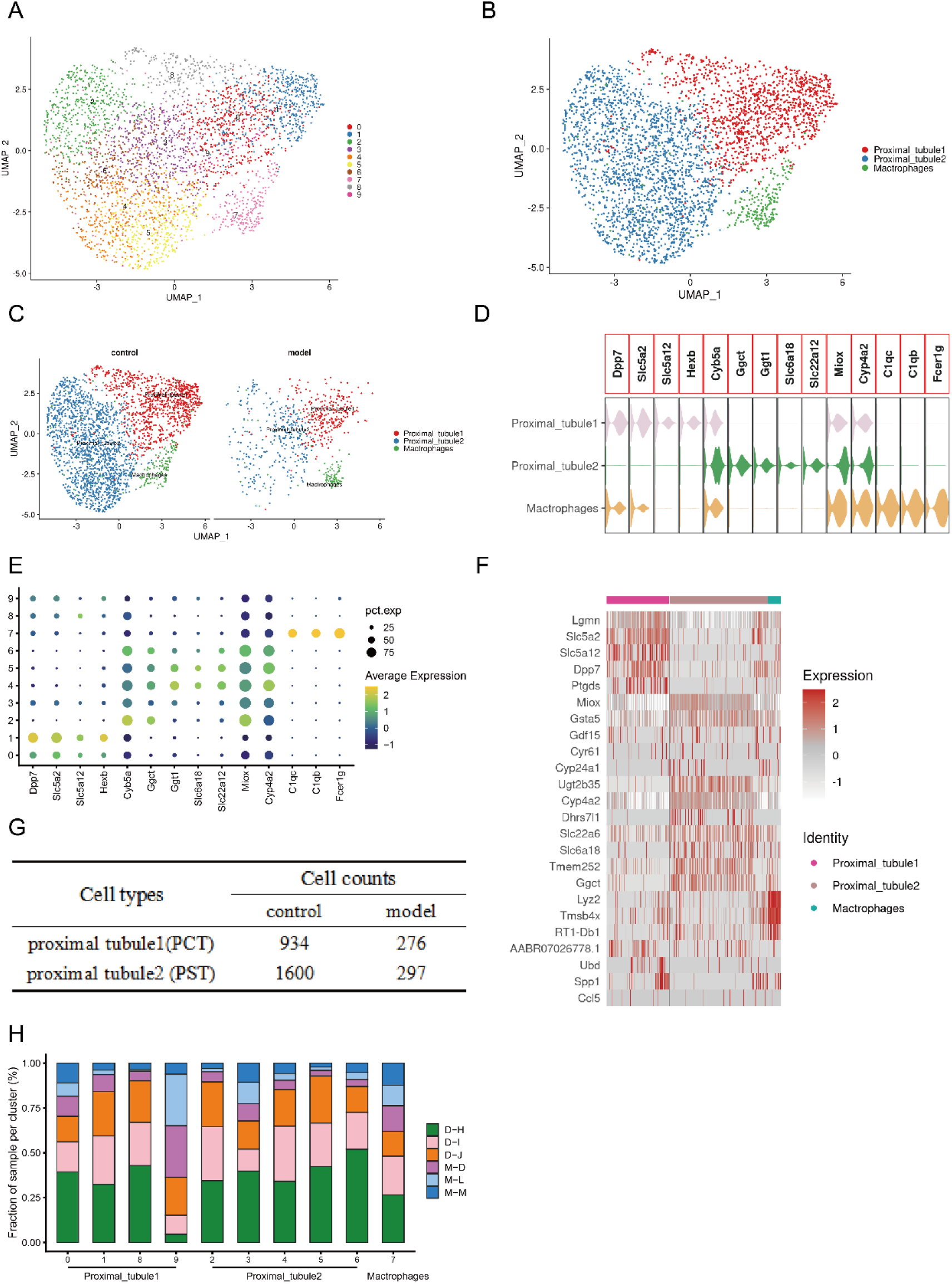
Proxmial tubule grouping. A, Re-clustering of PT using UMAP algorithm. B, PT cells are grouped according to the marker gene. C, UMAP visualization of heterogeneity of PT cells in the control and model groups. D, The violin diagram of marker gene. E, Scattered distribution of marker genes in PT cell typing. F, The marker gene heatmap of PT cell typing. G, Table shows the quantitative distribution of different subtypes of PT in the control and model group. H, The distribution plot of contribution of cells to different subtypes of PT in different samples.

Using Monocle 3.0 software for pseudotime analysis, PT cell development was divided into 9 states. With the growth of the pseudotime sequence, PT cells gradually differentiated from macrophages, PCT cells to PST cells (Figure 7A). Cd74, Fcer1g, Lyz2, Tyrobp were expressed more at the early stage of differentiation, and with the increase of the pseudotime sequence, the relative expression decreased, and the relative expression of Gsta1 and Rida increased (Figure 7B-7C). The relative genes expression heatmap of genes in cell fate branches is shown in Figure 7D-E.

**Fig. 7.**
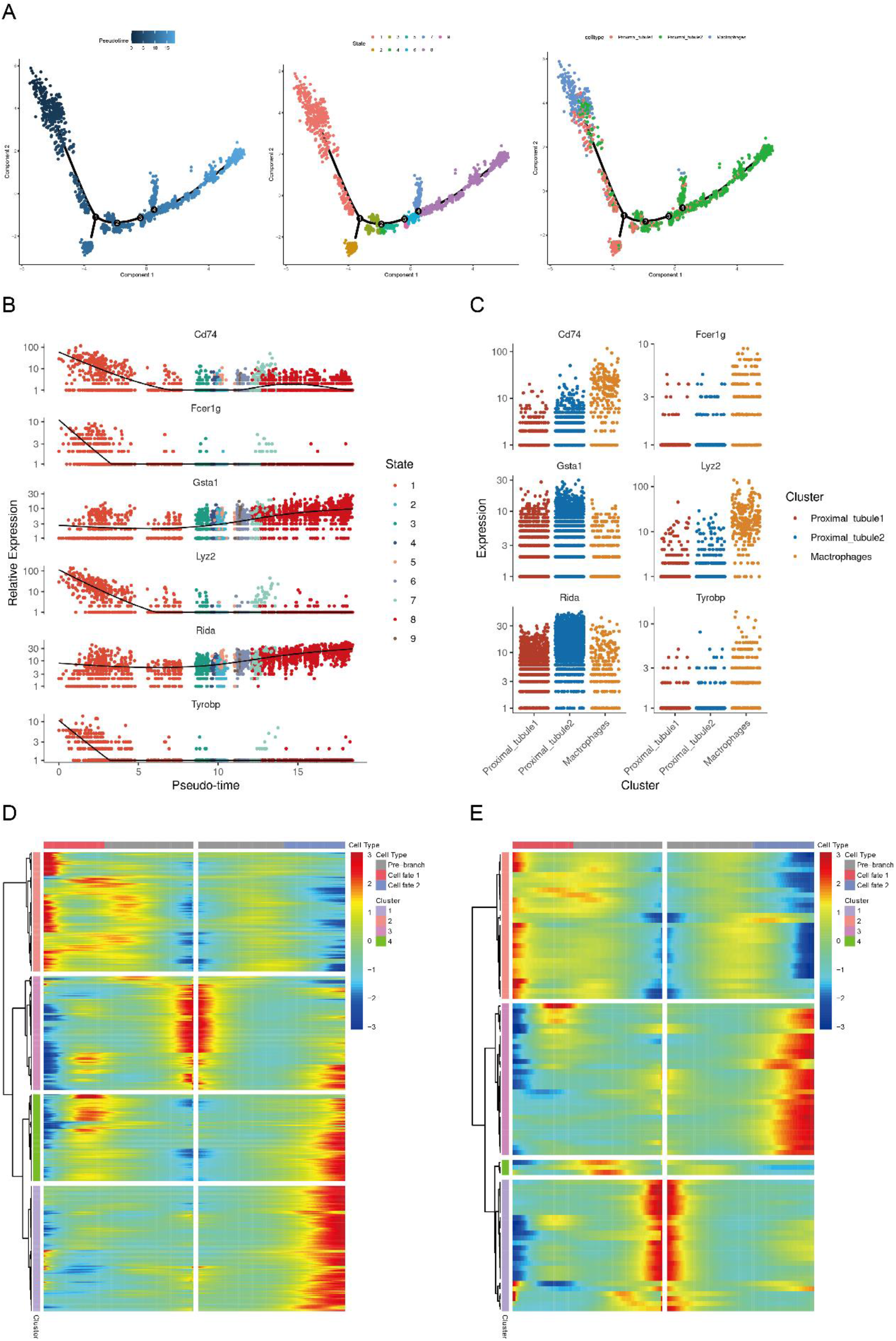
Evolution of PT differentiation. A, The reference timeline of PT cell development. B, The relative expression of genes in PT cell developmental trajectories. C, The relative expression of genes of different PT cell subsets in developmental trajectories. D, Gene expression heatmap of cell branch 1. E, Gene expression heatmap of cell branch 2.

### Function enrichment analysis of PT cell

The differential expression and enrichment analysis of genes in the control and model groups in the PT cell population are shown in Figure 8. Compared with the control group, a total of 148 genes were upregulated and 91genes were downregulated in the model group (Figure 8A). As shown in Figure 8B-E, there were 206 differential genes (71 up-regulated and 135 down-regulated) between PST and PCT in the control group, 137 differential genes (36 up-regulated, 101 down-regulated) between PST and PCT in the model group, 185 differential genes (63 up-regulated, 122 down-regulated) between the control and model groups in the PCT cell population, and 261 differential genes (96 up-regulated, 165 down-regulated) between the control and model groups in the PST cell population. GO enrichment of differential genes in PT cell populations between control and model groups showed that the upregulated genes in the control group were mainly related to fatty acid metabolic process, cellular amino acid metabolic process and generation of precursor metabolites and energy (Figure 8F). The genes downregulated by PCT compared with PST in the control group and model group involved BP such as fatty acid metabolic process and glutathione metabolic process, and were also related to pathways such as glutathione metabolism and fatty acid degradation, and downregulated genes were enriched in PPAR signaling pathway (Supplementary fig. 1, Supplementary fig. 2). In PT cell populations (PCT and PST), the genes upregulated in the control group compared to the model group involved fatty acid metabolic processes, and were also associated with pathways such as glutathione metabolism and fatty acid degradation (Supplementary fig. 3, Supplementary fig. 4). KEGG pathway analysis showed that upregulated genes in the control group were involved in fatty acid degradation (Figure 8G). The genes in the control group were mainly involved in BP such as negative regulation of immune system process, leukocyte migration, and cell chemotaxis. The CC are mainly involved in lysosome and lytic vacuole (Figure 8H). The downregulated genes are mainly related to pathways such as antigen processing and presentation, lysosome, and apoptosis (Figure 8I).

**Fig. 8.**
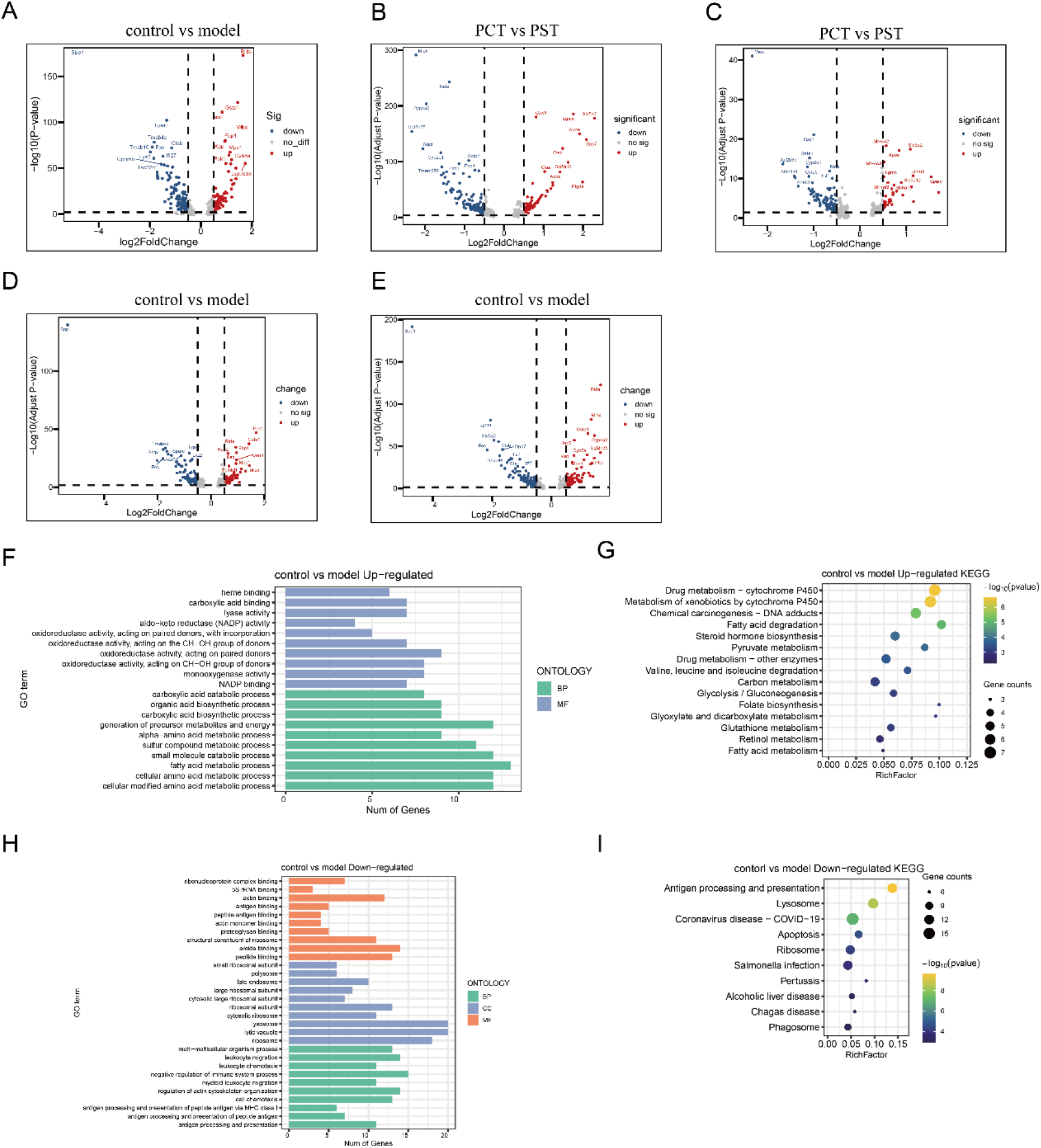
After PT was re-clustered, the differential gene enrichment analysis between control and model group was analyzed. A, Volcano plot of differential genes between control and model groups. B, Volcano plot of the difference gene between PCT and PST in the control group. C, Volcano plot of the difference gene between PCT and PST in the model group. D, Volcano plot of differential genes of PCT cell population in control and model groups. E, Volcano map of differential genes of PST cell population in control and model groups. F, GO enrichment of differential genes upregulated in the control group compared to the model group. G, KEGG pathway enrichment of differential genes upregulated by the control group compared to the model. H, GO enrichment of differential genes downregulated by the control group compared to the model. I, Enrichment of KEGG pathway of differential genes downregulated by the control group compared with the model.

### HAO2 promotes fatty acid metabolic process

In the functional enrichment analysis of PT cells, fatty acid metabolic process-related genes were found to be downregulated in the model group, where HAO2 is the gene involved in regulating fatty acid metabolic process. To further validate the role of HAO2 and fatty acid metabolic processes in CKD. Immunofluorescence was used to verify HAO2 expression in kidney tissue (Figure 9A), oil red O staining was used for lipid accumulation detection (Figure 9B), and ATP (Figure 9C) and FFA (Figure 9D) content was detected using ELISA. These results showed that HAO2 expression was localized in the renal tubules, and the model group was significantly reduced compared with the control group. Compared with the control group, the accumulation of lipid droplets increased in the model group, the FFA was significantly increased in the model group, and ATP was depleted.

**Fig. 9.**
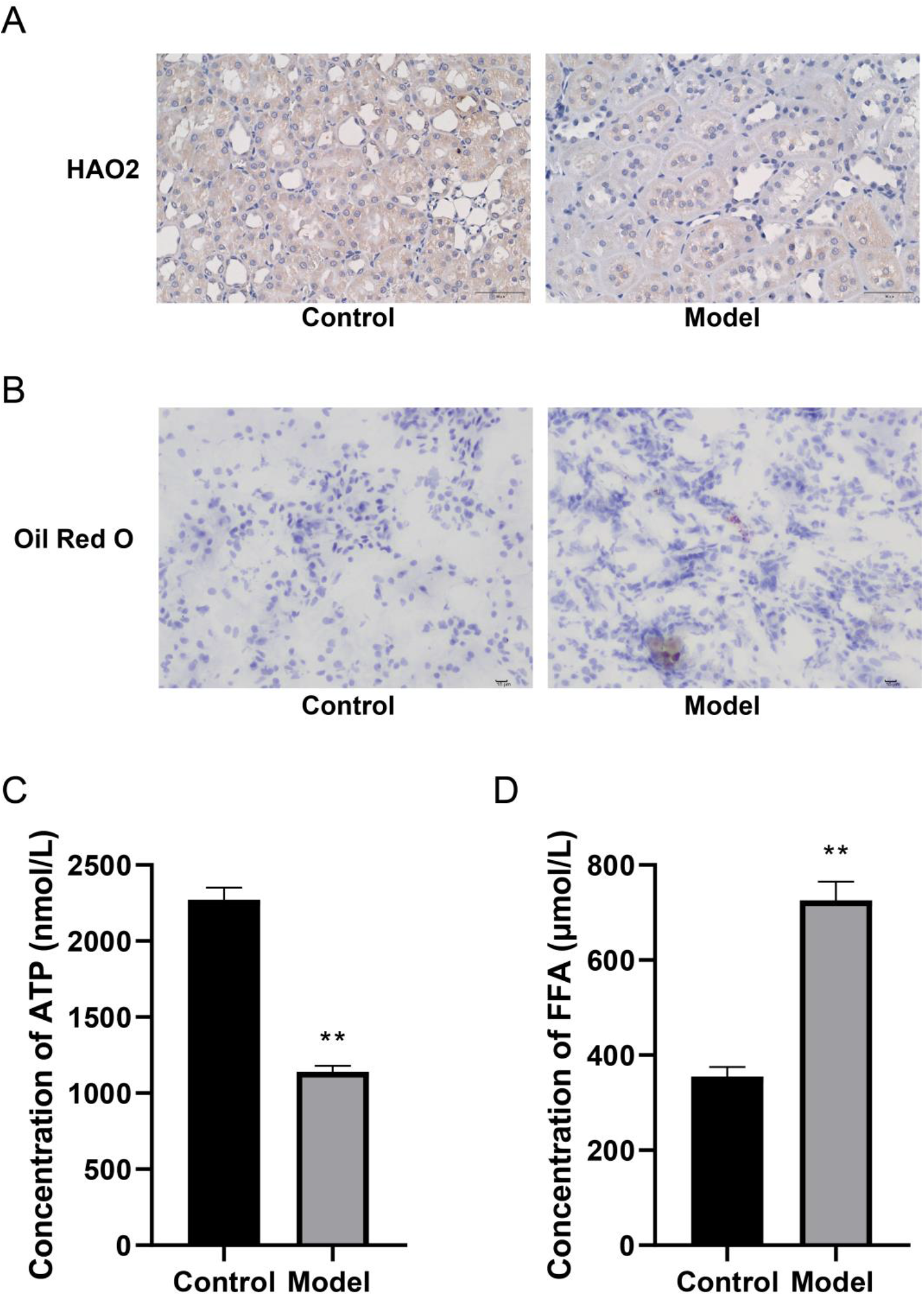
Detection of lipid accumulation in control and model group. A. Immunofluorescence is used to verify the expression of HAO2 in renal tissue. x400. B. Oil red O staining for lipid accumulation detection. x400. C, ELISA detects ATP and D, FFA content. *P<0.05, the difference between the two groups is significant; **P<0.01, the difference between the two groups is very significant.

### Fatty acid oxidation is inhibited in CKD

To further investigate the key role of HAO2 and fatty acid metabolism in CKD. We used qPCR and WB to detect ATP receptor levels and expression of key proteins for fatty acid oxidation. The results are shown in Figure 10, the expression of ATP receptor AMPK, ACC gene and protein did not differ significantly in the control and model group, while the phosphorylation level of AMPK, ACC protein increased significantly in the model group. The gene and protein expression levels of the key fatty acid oxidation factors ACOX1, PPARα and PGC1α were significantly reduced in the model group, indicating that the fatty acid oxidation pathway was inhibited in CKD.

**Fig. 10.**
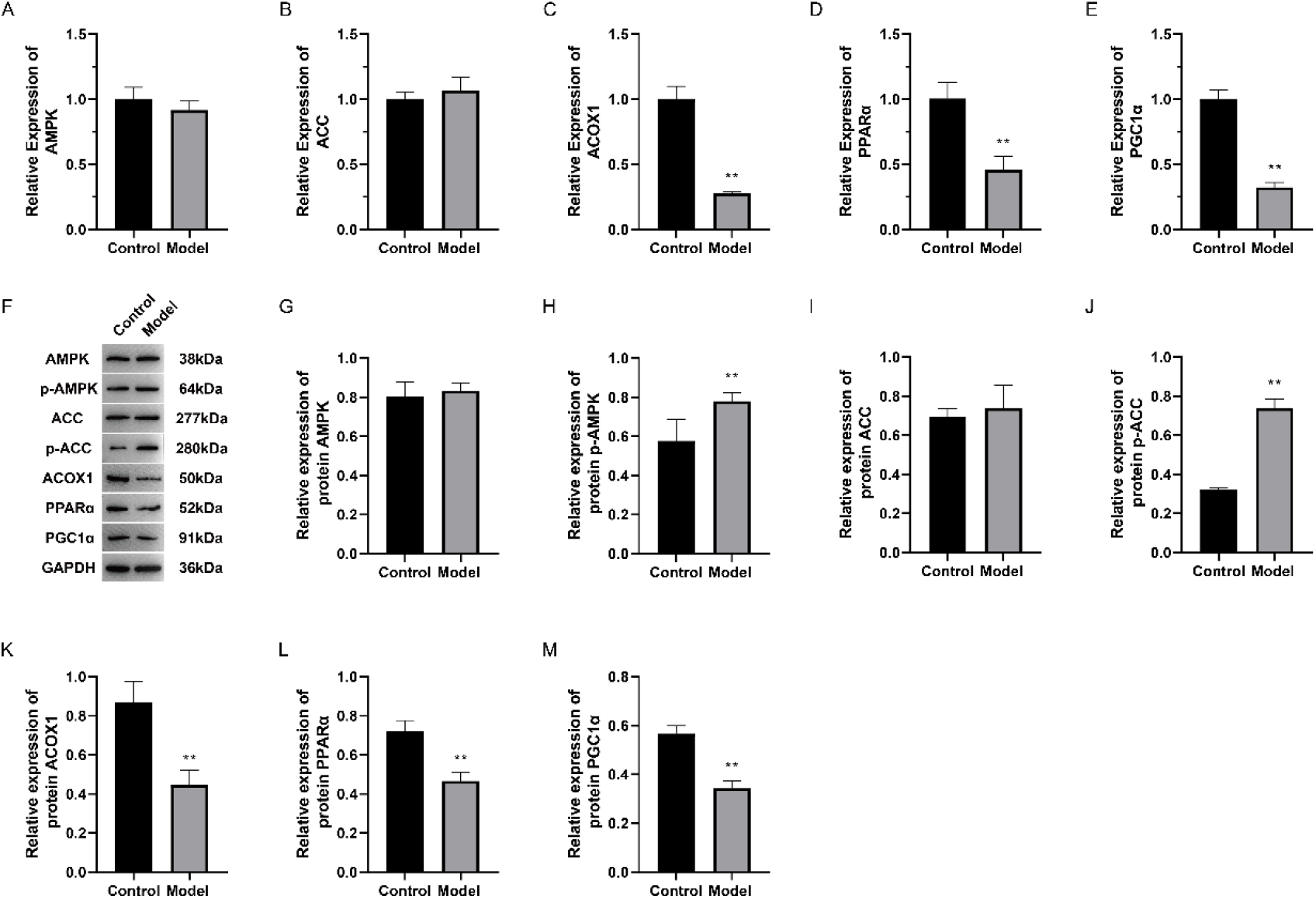
Fatty acid oxidation is inhibited in kidney injury. qPCR used for verification of A, AMPK; B, ACC; C, ACOX1; D, PPARα and E, PGC1α gene transcription levels. F, WB immunoblotting representative blots. G, AMPK; H, p-AMPK; I, ACC; J, p-ACC; K, ACOX1; L, PPARα and M, PGC1α proteins are relatively quantitative. *P<0.05, the difference between the two groups is significant; **P<0.01, the difference between the two groups is very significant.

### Overexpression of HAO2 enhances fatty acid metabolism by promoting fatty acid oxidation pathways

In order to further clarify the key role of HAO2 in fatty acid metabolism, the effect on fatty acid metabolism was studied by overexpressing HAO2 and interfering with HAO2 gene. Compared to the control group, lipid droplet accumulation decreased in the HAO2 overexpression group, while increased after interfering with HAO2 expression (Figure 11A). ATP content decreased significantly in the HAO2 interference group and increased significantly in the HAO2 overexpression group (Figure 11B). Similar to the lipid droplet accumulation results, FFA decreased in the HAO2 overexpression group, while FFA increased after interfering with HAO2 expression (Figure 11C).

HAO2 mRNA increased and decreased, respectively, in the HAO2 overexpression and interference groups compared to the control group (Figure 12A). After overexpression or interference with HAO2, there was no significant difference in AMPK and ACC gene expression (Figure 12B, D). ACOX1, PPARα and PGC1α mRNAs increased and decreased significantly in the HAO2 overexpression and interference groups, respectively (Figure 12C, E-F).

**Fig. 11.**
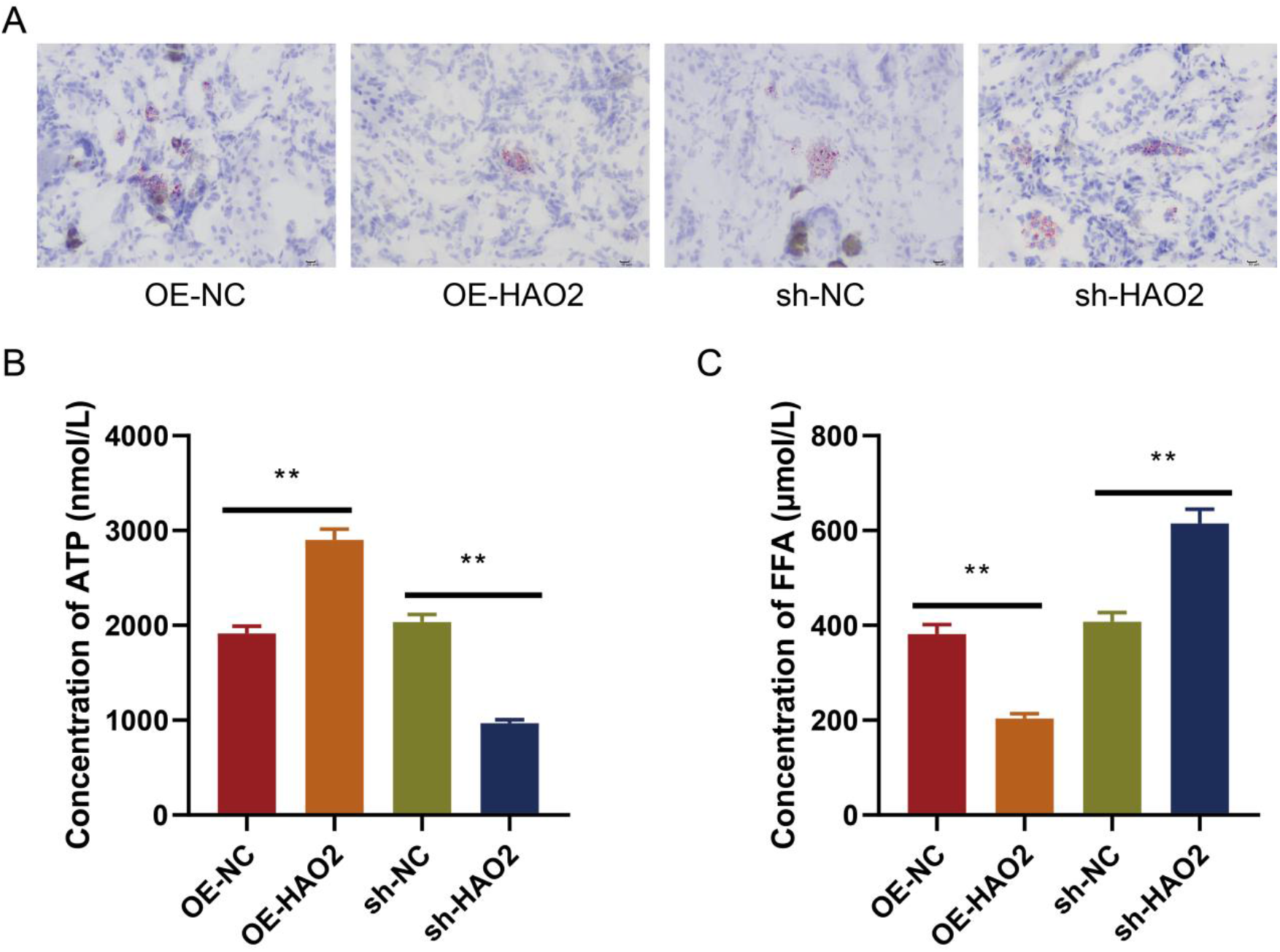
Overexpression of HAO2 reduces lipid accumulation. A, Oil red O staining for lipid accumulation detection. x400. B, ELISA detects ATP and C, FFA content. *P<0.05, the difference between the two groups is significant; **P<0.01, the difference between the two groups is very significant.

**Fig. 12.**
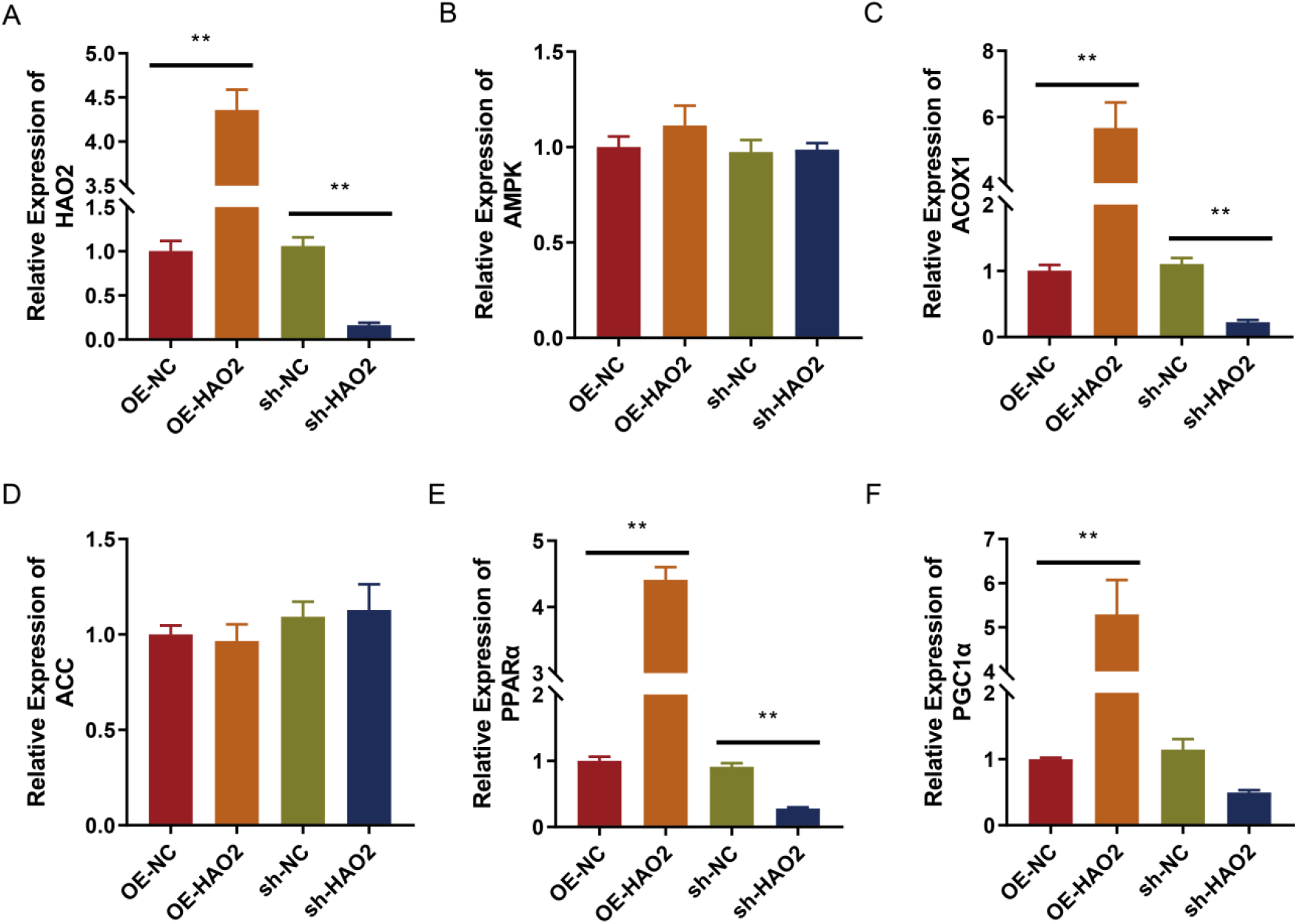
Overexpression of HAO2 promotes fatty acid oxidation pathways. qPCR used for verification of A, HAO2; B, AMPK; C, ACOX1; D, ACC; E, PPARα and F, PGC1α gene levels. *P<0.05, the difference between the two groups is significant; **P<0.01, the difference between the two groups is very significant.

The expression of these genes at the protein level by WB (Figure 13A) showed that HAO2 protein increased and decreased significantly in the HAO2 overexpression and interference groups, respectively (Figure 13I), similar to the qPCR results. AMPK, ACC protein after overexpression or interference with the HAO2 gene was not significantly different (Figure 13B, D). However, AMPK, ACC protein phosphorylation levels are significantly reduced and elevated after overexpression or interference with the HAO2 gene, respectively (Figure 13C, E). ACOX1, PPARα, and PGC1α proteins increased and decreased significantly in the HAO2 overexpression and interference groups, respectively (Figure 13F-H).

**Fig. 13.**
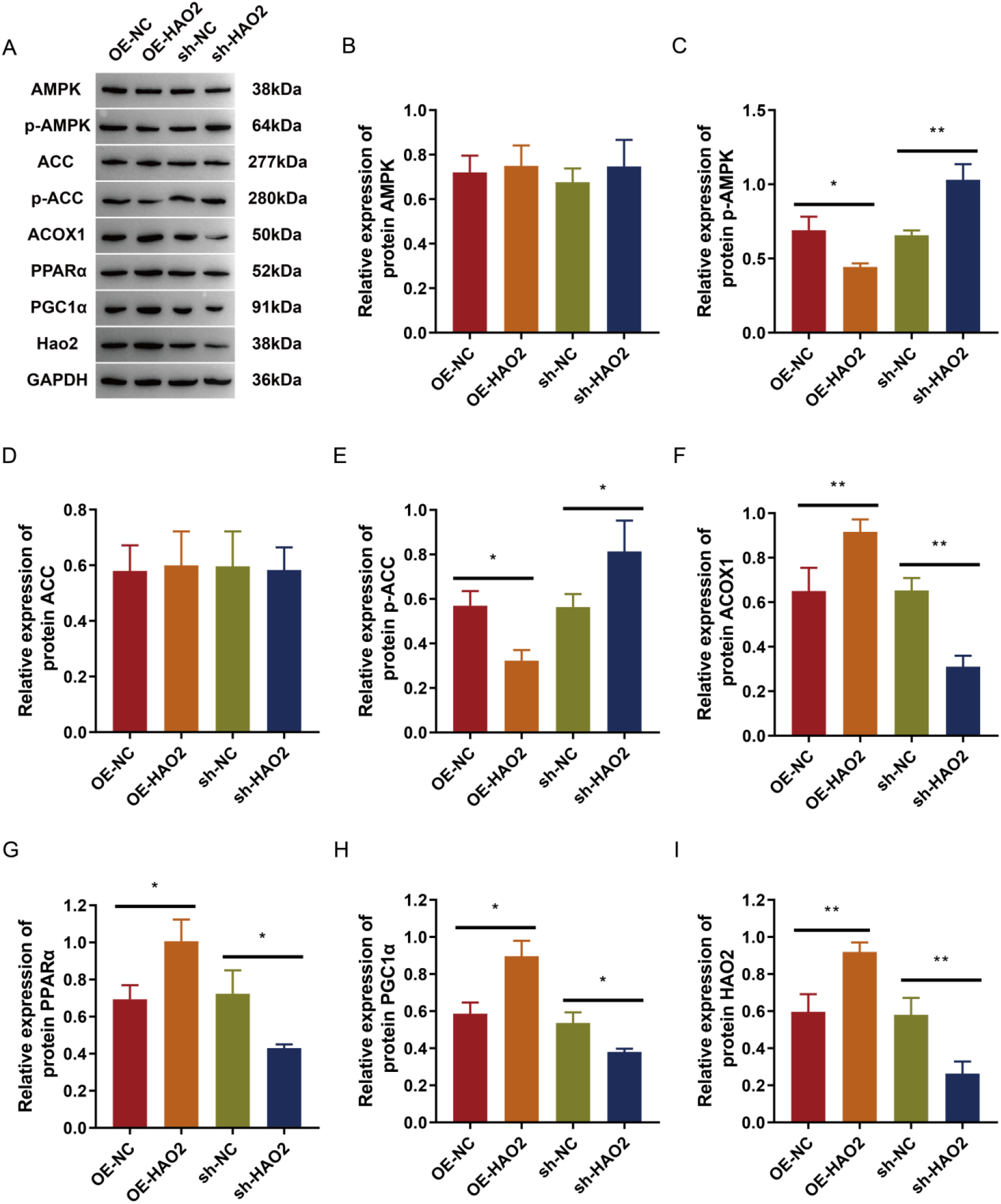
WB was used to detect relevant protein expression. A, WB immunoblotting representative blots. B, AMPK; C, p-AMPK; D, ACC; E, p-ACC; F, ACOX1; G, PPARα; H, PGC1α and I, HAO2 proteins are relatively quantified. *P<0.05, the difference between the two groups is significant; **P<0.01, the difference between the two groups is very significant.

## Discussion

Fatty acids can be divided into short-chain fatty acids (carbon atom number less than 6), medium-chain fatty acids (carbon atom number between 7-12) and long-chain fatty acids (carbon chain length greater than 12) according to the length of the carbon chain, of which long-chain fatty acids are the main source of energy synthesis, fatty acid oxidation can be used as the main energy source of some cells with strong metabolic function to produce more ATP than glucose. At the same time, long-chain fatty acids are also the main components of membranes and lipids. In 1982, Moorhead first proposed the “lipid-nephrotoxicity hypothesis”, which holds that in kidney disease, the liver produces increased lipoprotein synthesis, forming lipid-mediated damage cycles, and glomerular and tubulointerstitial damage worsens[13]. Then this hypothesis has been confirmed by a series of clinical and animal studies and is generally accepted. The adult rats fed with the addition of 3-4% cholesterol can lead to arterial damage and glomerular abnormalities, glomerulosclerosis and abnormalities in rat kidney tissue, and lipid droplets are found [14, 15]. The rat adenine diet model has also been found to cause rapid-onset nephropathy tubulointerstitial fibrosis, tubular atrophy, crystal formation, and marked vascular calcification. Lower intake of adenine results in slowly progressive renal impairment cardiovascular disease, allowing for relatively stable renal and cardiovascular disease, similar to CKD in humans [12].

Excessive accumulation of triglycerides can lead to cytolipid toxicity that may lead to the development of fibrosis, and sebum accumulation on tubules has received widespread attention in the setting of acute and diabetic nephropathy[16, 17]. Expression of key enzymes and regulators of fatty acid oxidation (FAO) is lower in human and mouse models of tubulointerstitial fibrosis, while lipid deposition increases intracellular [18]. Fatty acids can be degraded in mitochondria by β-oxidation pathway. Free fatty acids can be absorbed by the liver and converted to triglycerides, and lipid abnormalities have been demonstrated in ccRCC, including cholesterol and triglycerides[19]. Inhibition of fatty acid oxidation by epithelial cells in tubules leads to ATP depletion, intracellular lipid deposition, and observation of fibrotic phenotypes [18].

In this study, we performed scRNA-seq by collecting blood samples and kidney tissue from normal control rats and rats with CKD, analyzed the expression profiles of 27,332 cells, and plotted single-cell atlas. The results showed that the cells were mainly clustered into 23 cell subsets, mainly including 14 kinds of cells: MPCs, PT, Tc, DCT, B-IC, A-IC, CNT, ALOH, BC, Neu, Endo, Pla, NKT, Baso, among which the number of NKT and Baso was small, and the number of MPCs, PT and Tc was greater. In addition, MPCs, Neu increased in the model group, and PT cells and Tc decreased in the model group, suggesting that MPCs, PT cells and Tc played an important role in the process of CKD. We further re-cluster cell subsets of MPCs and PT cells with a large number of cells and strong heterogeneity. The regrouping results of MPCs showed that MPCs cells had 15 cell subsets, mainly containing macrophages and monocytes. Renal mononuclear phagocytes, such as dendritic cells and macrophages, express NLRP3 inflammatory body components and may induce cell death by activating caspase-1. Lipid-mediated mononuclear phagocytes infiltrate can engulf lipid-forming foam cells, releasing multiple cytokines and proteases leading to glomerular damage.

The subpopulation of PT cells divides PT cells into two cell subtypes: PCT and PST. GO and KEGG enrichment analysis of PT cell populations showed that the genes upregulated in the control group were mainly related to fatty acid metabolic process and fatty acid degradation, and the genes downregulated by PCT compared with PST in the control group and model group were mainly enriched in fatty acid metabolic process and fatty acid degradation. It suggests that PST subtypes in PT cells may play an important role in promoting fatty acid metabolic process and fatty acid degradation, and promoting or restoring fatty acid metabolism may be beneficial in the treatment of CKD in rats. In patients with CKD, excess fatty acids are deposited in the proximal tubules of the kidneys, and proteinuria develops. In the physiological state of CKD, albumin binds to free fatty acids after filtration from the glomerulus, and then is reabsorbed by the PT, releasing fatty acids to liver-type fatty acid-binding protein (L-FABP), and transporting to lysosomes to bind to fatty acids, and L-FABP is redistributed to the cell matrix peroxisome, participating in the peroxidation reaction [20]. Fatty acids that are not oxidized after peroxidation are cytotoxic and can cause inflammatory factor production, macrophage infiltration, and accelerate tubulointerstitial damage [21, 22].

From the results of GO and KEGG enrichment analysis of the PT cell population in Figure 8, the fatty acid metabolic process was downregulated in the model group, and the HAO2 gene was significantly downregulated in the model group. In order to further verify the role of HAO2 gene in CKD, we first determined by immunohistochemistry experiments that HAO2 was localized in the renal tubules, and the model group was significantly reduced compared with the control group, the ATP content was reduced in the model group, and the oil red O staining indicated that lipid droplets increased in the model group, and lipid droplets decreased in tissues after overexpression of HAO2, suggesting that HAO2 may be involved in promoting fatty acid metabolism. The results of protein detection showed that AMPK, ACC phosphorylated protein were significantly elevated in the model group, and reduced after overexpression of HAO2, key factors of fatty acid oxidation such as ACOX1, PPARα and PGC1α decreased in the model group, and after overexpression of HAO2, the levels of ACOX1, PPARα, PGC1α and other genes and phosphorylated proteins increased. Therefore, HAO2 may be an important regulator of fatty acid metabolism in CKD, and overexpression of HAO2 can enhance fatty acid metabolism by promoting fatty acid oxidation pathway.

The HAO2 gene belongs to the hydroxy acid oxidase family and encodes a peroxisome protein with 2-hydroxyacid oxidase activity, which is downregulated in hepatocellular carcinoma (HCC), overexpressing HAO2 can impair the growth of HCC cells by increasing reactive oxygen species (ROS) production and lipid peroxidation[23]. Wen Xiao et al. reported in 2019 that HAO2 can inhibit the malignant tumor process of renal clear cell carcinoma, and the mechanism to inhibit malignant tumors may be promoting lipid metabolism [24].

In summary, this study constructed a rat model of CKD, drew a single-cell atlas of CKD in rats, found cell subsets related to the process of CKD, described the cell evolution trajectory and functional enrichment of related cell subsets, and discovered the possible key role of fatty acid metabolism processes in CKD. Through *in vitro* verification experiments, it was found that HAO2 may be an important regulator of fatty acid metabolic process in CKD, and overexpression of HAO2 can enhance fatty acid metabolism by promoting fatty acid oxidation pathway.

## Data Availability Statement

The data used to support the findings of this study are included within the article.

