## Supplementary figures for "Role of HAO2 in rats with chronic kidney disease by regulating fatty acid metabolic processes in renal tissue"

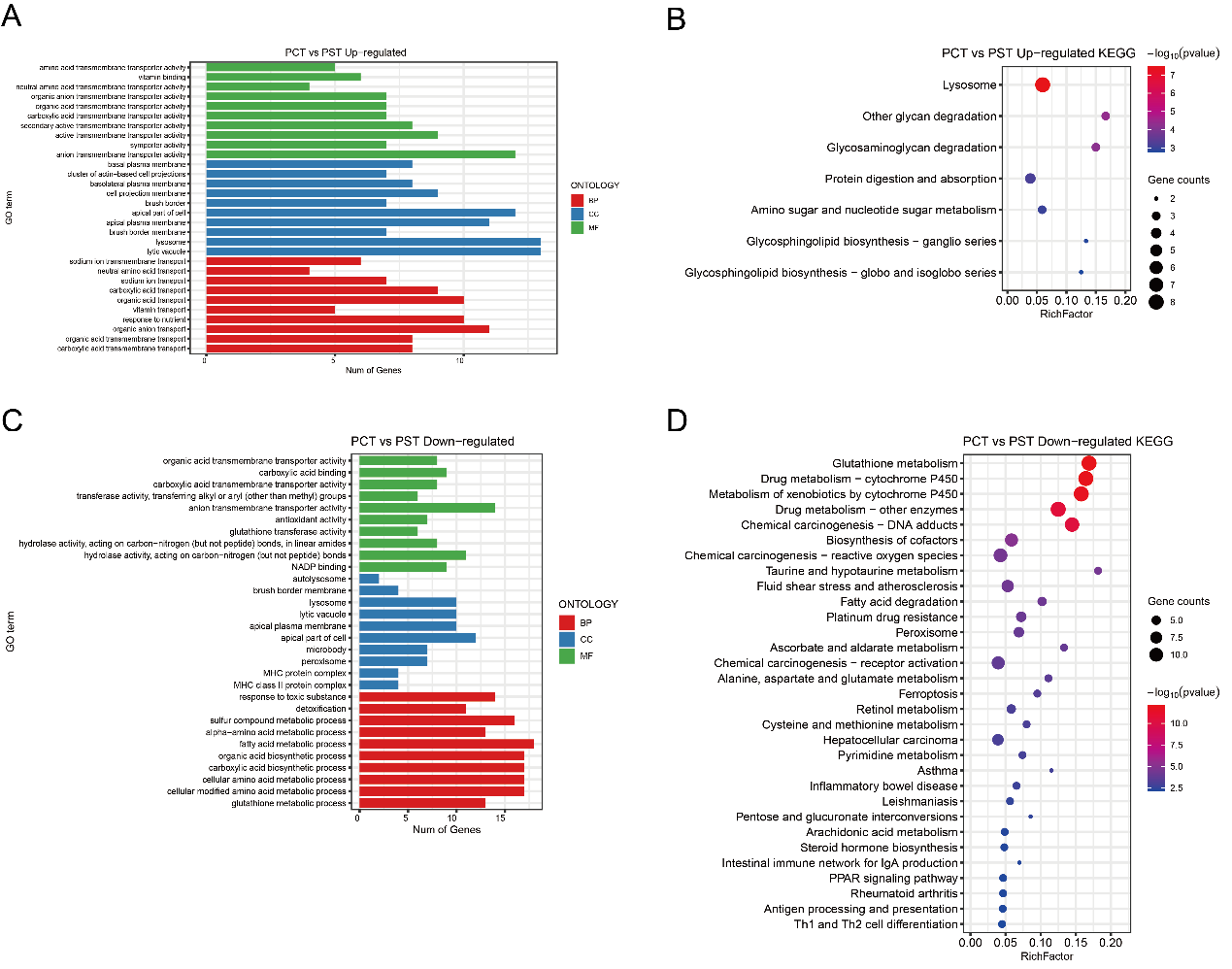

Supplementary fig. 1 Differential gene enrichment analysis between PT cell subtypes PCT and PST in the control group. A, GO enrichment of differential genes upregulated in PCT compared to PST in the control group. B, KEGG pathway enrichment of differential genes upregulated in PCT compared to PST in the control group. C, GO enrichment of differential genes downregulated in PCT compared to PST in the control group. D, KEGG pathway enrichment of differential genes downregulated in PCT compared to PST in the control group.

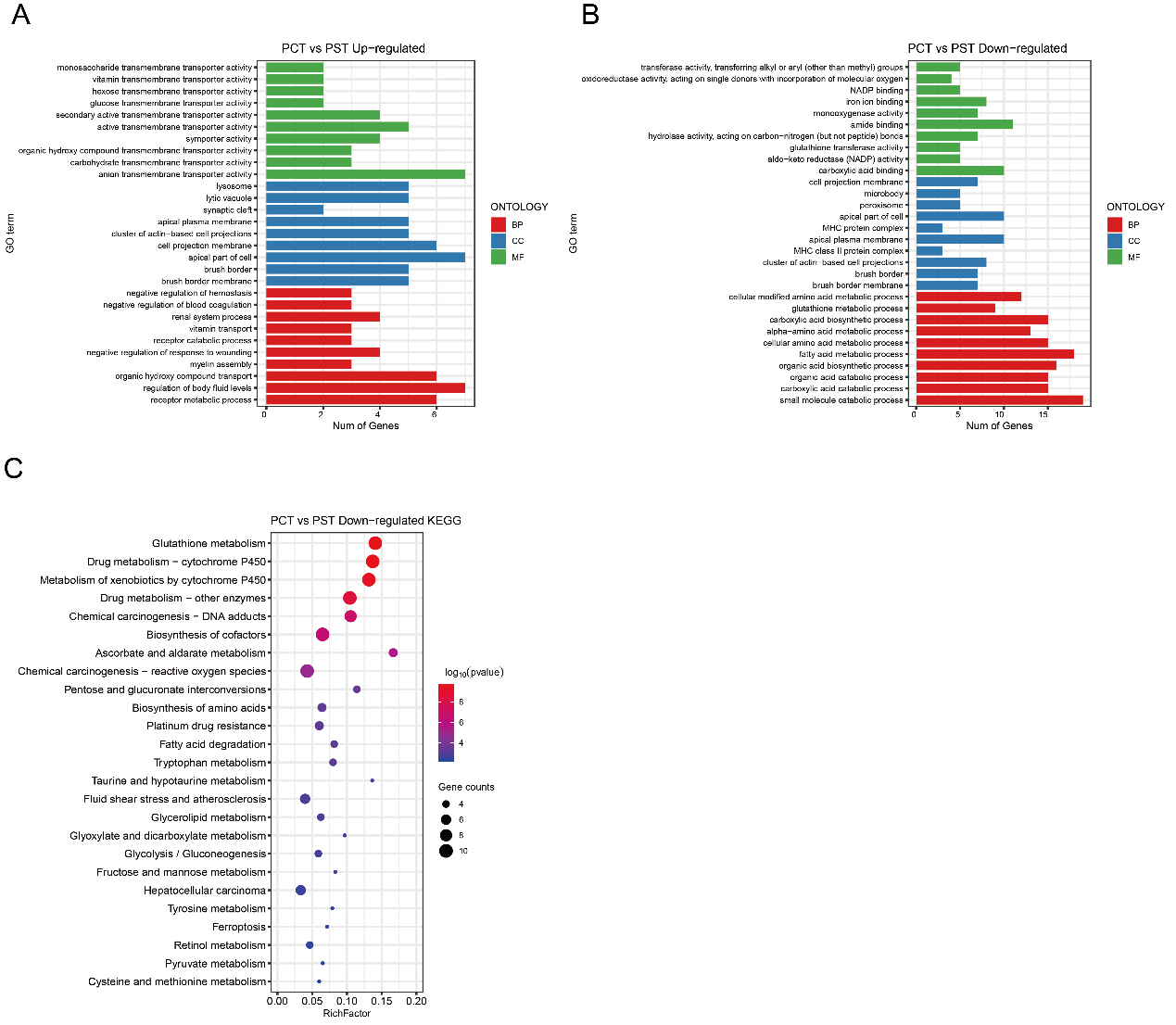

Supplementary fig. 2 Differential gene enrichment analysis between PT cell subtypes PCT and PST in the model group. A, GO enrichment of differential genes upregulated in PCT compared to PST in the model group. B, GO enrichment of differential genes downregulated in PCT compared to PST in the model group. C, KEGG pathway enrichment of differential genes downregulated in PCT compared to PST in the model group.

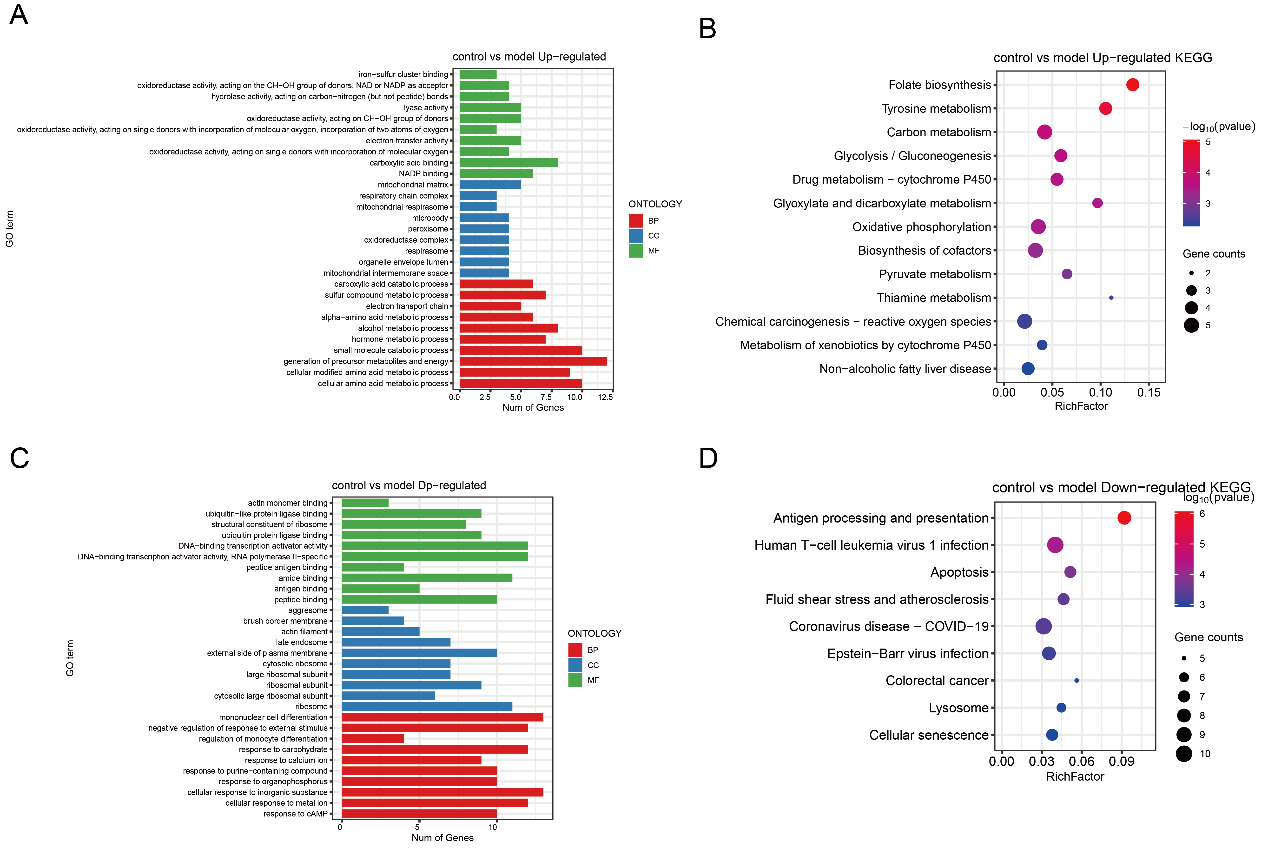

Supplementary fig. 3 Differential gene enrichment of PCT cells in control and model groups. A, GO enrichment of differential genes of PCT upregulated in control group compared to the model group. B, KEGG pathway enrichment of differential genes of PCT upregulated in control group compared to the model group. C, GO enrichment of differential genes of PCT downregulated in control group compared to the model group. D, KEGG pathway enrichment of differential genes of PCT downregulated in control group compared to the model group.

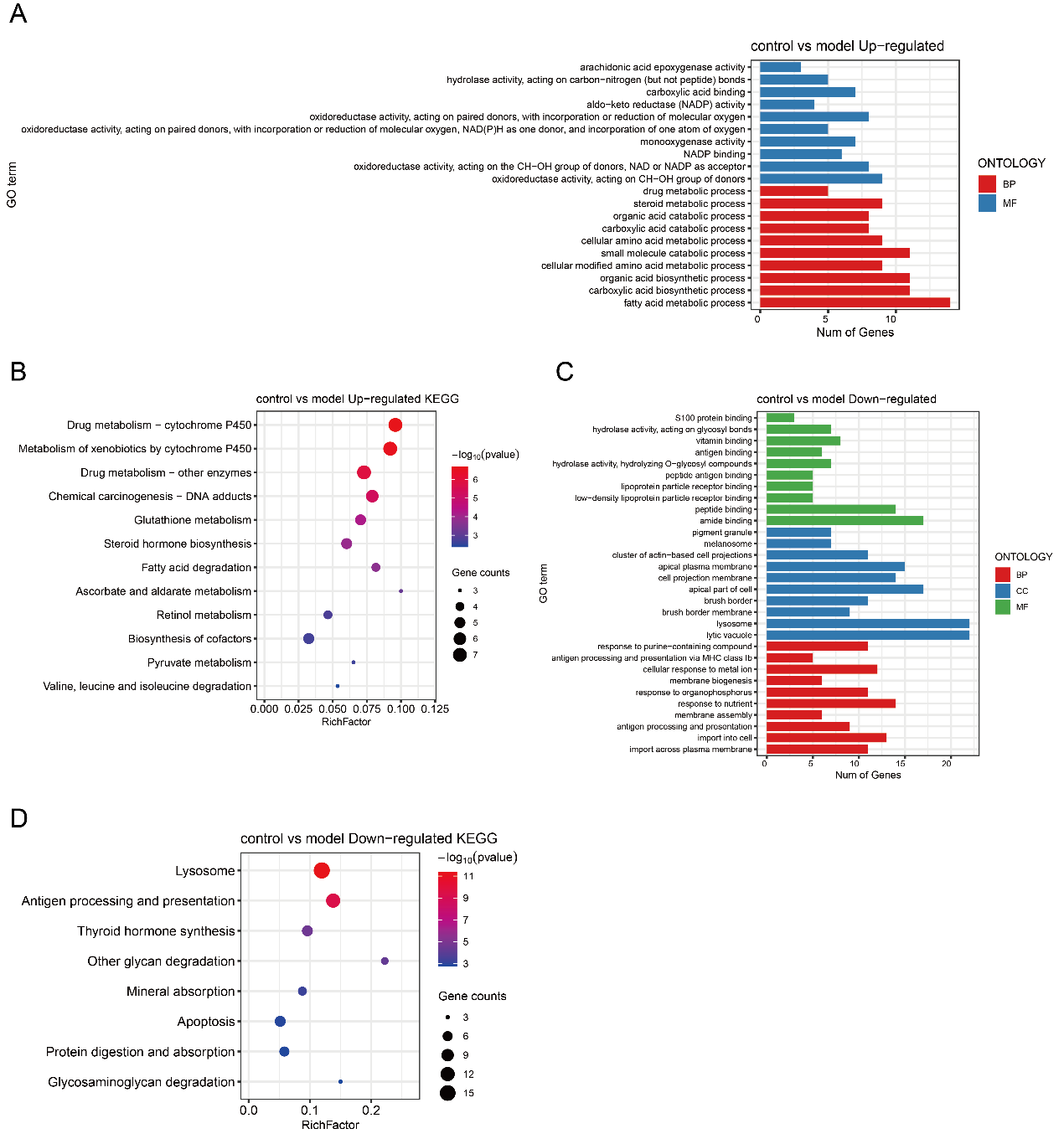

Supplementary fig. 4 Differential gene enrichment of PST cells in control and model groups. A, GO enrichment of differential genes of PST upregulated in control group compared to the model group. B, KEGG pathway enrichment of differential genes of PST upregulated in control group compared to the model group. C, GO enrichment of differential genes of PST downregulated in control group compared to the model group. D, KEGG pathway enrichment of differential genes of PST downregulated in control group compared to the model group.
